# Missing-link fold reveals the evolutionary pathway between RNA polymerase and ribosomal proteins

**DOI:** 10.1101/2023.07.05.547881

**Authors:** Sota Yagi, Shunsuke Tagami

## Abstract

Numerous molecular machines are required to drive the central dogma of molecular biology. However, the means by which such numerous proteins emerged in the early evolutionary stage of life remains enigmatic. Many of them possess small β-barrel folds with different topologies, represented by DPBB conserved in DNA and RNA polymerases, and RIFT, OB, and SH3 in ribosomal proteins. Here, we discovered that the previously reconstructed ancient DPBB sequence could also adopt a novel β-barrel fold named DZBB, which shares similarities with RIFT and OB. Indeed, DZBB could be transformed into them through simple engineering experiments. Furthermore, the OB designs could be converted into SH3 by circular-permutation. These results indicate that these β-barrels diversified quickly from a common ancestor at the beginning of the central dogma evolution.

---

The central dogma of molecular biology is governed by numerous molecular machines, including DNA polymerases, RNA polymerases, and ribosomes. Despite the detailed understanding of their regulation mechanisms, the evolutionary origins of such complex molecular machines remain obscure.

The evolutions of some pivotal proteins in the central dogma may have originated from the well conserved small β-barrels within their core regions(*1*). For example, the core domains of DNA polymerase D (PolD) from euryarchaea and all cellular RNA polymerases are composed of two homologous β-barrels with six strands, “double-psi β-barrels (DPBBs)”(*2*–*4*) (Fig. S1A). The ribosomal protein L3 (rL3) and several translation factors have a similar but topologically distinct six-stranded β-barrel, “RIFT”(*5, 6*) (Fig. S1B). The structures of DPBB and RIFT have two-fold pseudo symmetry, indicating they originated as shorter homo-dimeric peptides(*5, 7*). Five-stranded β-barrel folds such as “OB” and “SH3” are often found in other ribosomal proteins and translation factors(*8*–*10*) (Fig. S1C and D). Given that these β-barrel domains are highly conserved across all extant organisms and play critical roles in replication, transcription, and translation, it is hypothesized that they were among the earliest components of the primordial central dogma machinery(*1, 2, 11*).

These four β-barrels (DPBB, RIFT, OB, and SH3) are classified into different folds in the SCOP, CATH, and ECOD protein databases(*12*–*15*), as they have distinct topologies. Even so, the partial structure and sequence similarities between these folds have been detected. Over the last two decades, meticulous comparative analysis of sequence motifs and partial structures have independently suggested that the DPBB-RIFT, RIFT-OB, and OB-SH3 pairs diverged from a common ancestral protein(*5, 6, 9, 16*) (Fig.S1). Despite these efforts, no experimental evidence has been provided to demonstrate that such drastic fold transitions could occur via a feasible pathway, probably because of the huge sequence/structure diversity between modern proteins with the different folds, especially between the pseudo-dimeric ones (DPBB and RIFT) and the monomeric ones (OB and SH3). Therefore, an experimental reconstruction of the ancient evolutionary process between these β-barrels has been awaited to reveal the profound protein fold evolution before the establishment of the central dogma.

We previously reconstructed the evolutionary pathway of the DPBB fold, initiated through the homo-dimerization of a half-sized peptide with about 40 amino acids, followed by gene duplication and fusion(*7*). Furthermore, by reducing the amino acid repertoire of the peptide, we have created the homo-dimeric DPBB fold comprising only seven amino acid types (Ala, Gly, Asp, Glu, Val, Lys, and Arg, named mk2h_ΔMILPYS; Fig. 1A and B), which could have been synthesized by immature translation systems in early life(*7, 17*). In this study, by using this simplified DPBB peptide as the starting template, we experimentally reconstructed the evolutionary pathways between the various ancient β-barrel folds in the central-dogma machinery through an unexpected missing link.

**Fig. 1.**
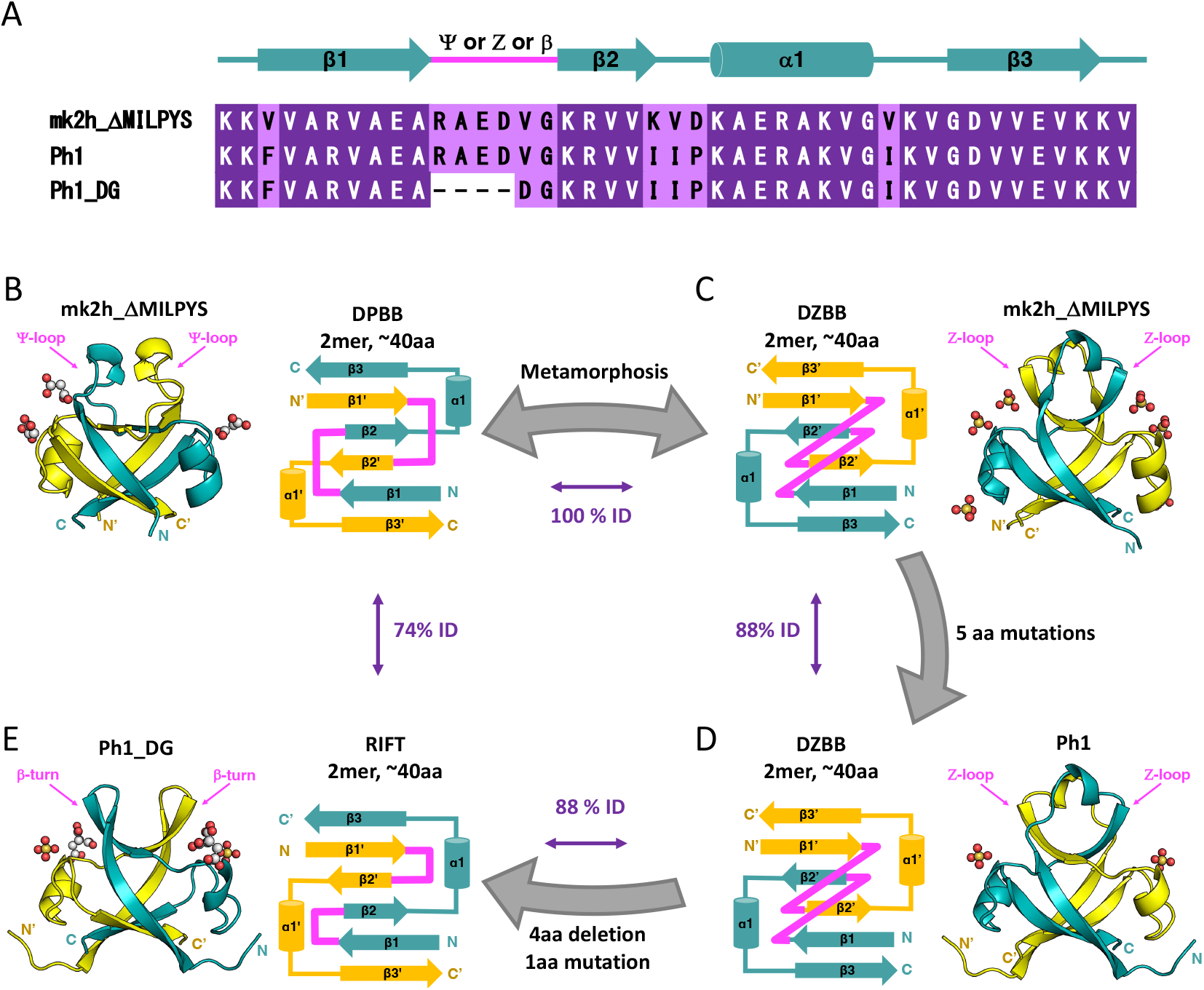
Experimental reconstruction of the evolutionary pathway from homo-dimeric DPBB to RIFT through a new barrel fold DZBB. (A) The amino acid sequences of the representative designs are aligned with the secondary structure elements. (B–E) The crystal structures and topological schemes of engineered proteins are shown. Single chains in each homo-dimeric structure are colored cyan and yellow. The 1st loops connecting β1-β2 are highlighted in pink in all panels. The sequence identities between the proteins are also shown between their panels. (B and C) mk2h_ΔMILPYS adopted in two folded states, DPBB and DZBB. (D) The five point mutations stabilized the DZBB fold in Ph1. (E) The remodeling of the loop connecting β1-β2 in Ph1 resulted in the RIFT-fold protein, PH1_DG.

## Conversion of homo-dimeric β-barrels through a missing-link fold

The most simplified DPBB protein we designed previously, mk2h_ΔMILPYS, did not fold in the typical buffer conditions (50 mM phosphate, 150 mM NaCl), but crystallized under two different conditions, containing malonate or malic acid ions, and adopted the DPBB fold in the crystals (Fig. 1A and B)(*7*). Here, we report its third type of crystal, formed under different conditions (100mM Tris, pH8.5, 20% PEG-400, 200mM lithium sulfate). Interestingly, we could not solve its structure by molecular replacement using the DPBB fold as a model, implying that mk2h_ΔMILPYS had adopted a different conformation in the third type of crystal.

At first, we expected that it adopted a RIFT-like structure, because DPBB and RIFT supposedly evolved from a common ancestral homo-dimeric peptide(*7*). Indeed, they commonly have (i) six-stranded β-barrel structures, (ii) an internal pseudo-two-fold symmetry, (iii) and a sequence motif “GD-box”, although the 1st loop configuration and the β2 direction are different (Fig. S2). Following this assumption, we tried to stabilize the unsolved conformation of mk2h_ΔMILPYS by introducing amino acid residues conserved in Phs018, a RIFT protein. Phs018 has high two-fold symmetry and likely retains the properties of the ancient RIFT-fold proteins(*5*). Phs018 has remarkable sequence similarity to mk2h_ΔMILPYS (20–27% identity; Fig S2), and we replaced five residues in mk2h_ΔMILPYS with to the ones at the corresponding positions of Phs018 (Fig. 1A). AlphaFold2 (AF2)(*18*) predicted that the resultant mutant, Ph1, would fold into a RIFT-like structure, albeit with a low lDDT region in the 1^st^ β-turn (Fig. S3). Ph1 was expressed, purified, and analyzed physiochemically. Circular dichroism (CD) and size exclusion chromatography (SEC) experiments indicated that Ph1 folded with β-rich characteristics and had moderate stability (Tm=58°C) (Fig. S4). Ph1 also formed crystals under similar buffer conditions to the third type of mk2h_ΔMILPYS crystal, including lithium sulfate.

By molecular replacement with the predicted Ph1 model and subsequent manual remodeling, the crystal structures of mk2h_ΔMILPYS (third type of crystal) and Ph1 were determined (Supplemental text 1). Surprisingly, they both adopted into a novel β-barrel fold that topologically differs from DPBB and RIFT (Fig. 1C and D). Compared to the homo-dimeric DPBB fold, the directions of the β2- and β2’-strands were inverted, resulting in all anti-parallel strand patterns like those in RIFT. However, the 1st loop connecting β1– β2 was rolled up, unlike the simple β-turn in RIFT (Fig. 1C, D, and S5A). As this loop configuration resembles the letter “Zeta” in the topological scheme, we named this new β-barrel fold the Double-Zeta β-barrel (DZBB). Therefore, mk2h_ΔMILPYS can fold into two different structures with the identical sequence, as a metamorphic protein. The five point-mutations derived from Phs018 (RIFT) in Ph1 stabilized the DZBB fold.

To further convert DZBB into RIFT, the sequence forming the Z-loop in Ph1 was replaced with two residues (DG, GD, or GG) facilitating β-turn formation(*19*) (Ph1_DG, Ph1_GD, and Ph1_GG) (Fig. S6 and Table S1). CD experiments demonstrated that Ph1_DG and Ph1_GD folded (Fig. S7), and Ph1_DG formed well-diffracting crystals. In contrast, Ph1_GG was unfolded but still formed crystals in the presence of sulfate ions. The crystallographic analysis revealed that Ph1_DG and Ph1_GG adopted the homo-dimeric RIFT-fold (Figs. 1E and S5B). Thus, the short In/Del at the 1st loop position is the determinant for the fold transition between DZBB and RIFT (Fig. 1A). The successful experimental conversion from DPBB to RIFT through the DZBB fold, by just a few mutations, indicates that DZBB is a missing link between the ancient homo-dimeric β-barrels in transcriptional and translational proteins (Fig. S1).

## Dual-folding of mk2h_ΔMILPYS induced by small ligands

mk2h_ΔMILPYS was originally obtained by substituting three tyrosine residues in the stable parent DPBB protein, mk2h_ΔMILPS, composed of eight amino acid types(*7*). Thus, these three mutations destabilized the DPBB fold and then allowed it to fold into the DZBB fold, resulting in the dual-folding property. Furthermore, the addition of five-point mutations in Ph1 stabilized the DZBB fold and abolished the capability to adopt the DPBB fold. Such metamorphic states like those of mk2h_ΔMILPYS could have existed to bridge between different folds smoothly during drastic fold transitions(*20, 21*).

Different ligand molecules probably induced the metamorphism in mk2h_ΔMILPYS. In the crystal structure of the DPBB-fold, two malonate ions bind to a positively charged pocket around the α1 helix, and are coordinated by Lys24, Arg27, and Lys33 (Fig. 2A). Alternatively, in the crystal structure of mk2h_ΔMILPYS with the DZBB fold, two additional residues, Arg21, and Arg18’ from the other chain, coordinate the two sulfates (Fig. 2B). The sulfate ions may attract these two additional residues in the folding process, and then stabilize the inverted β2-strand in the DZBB-fold.

**Fig. 2.**
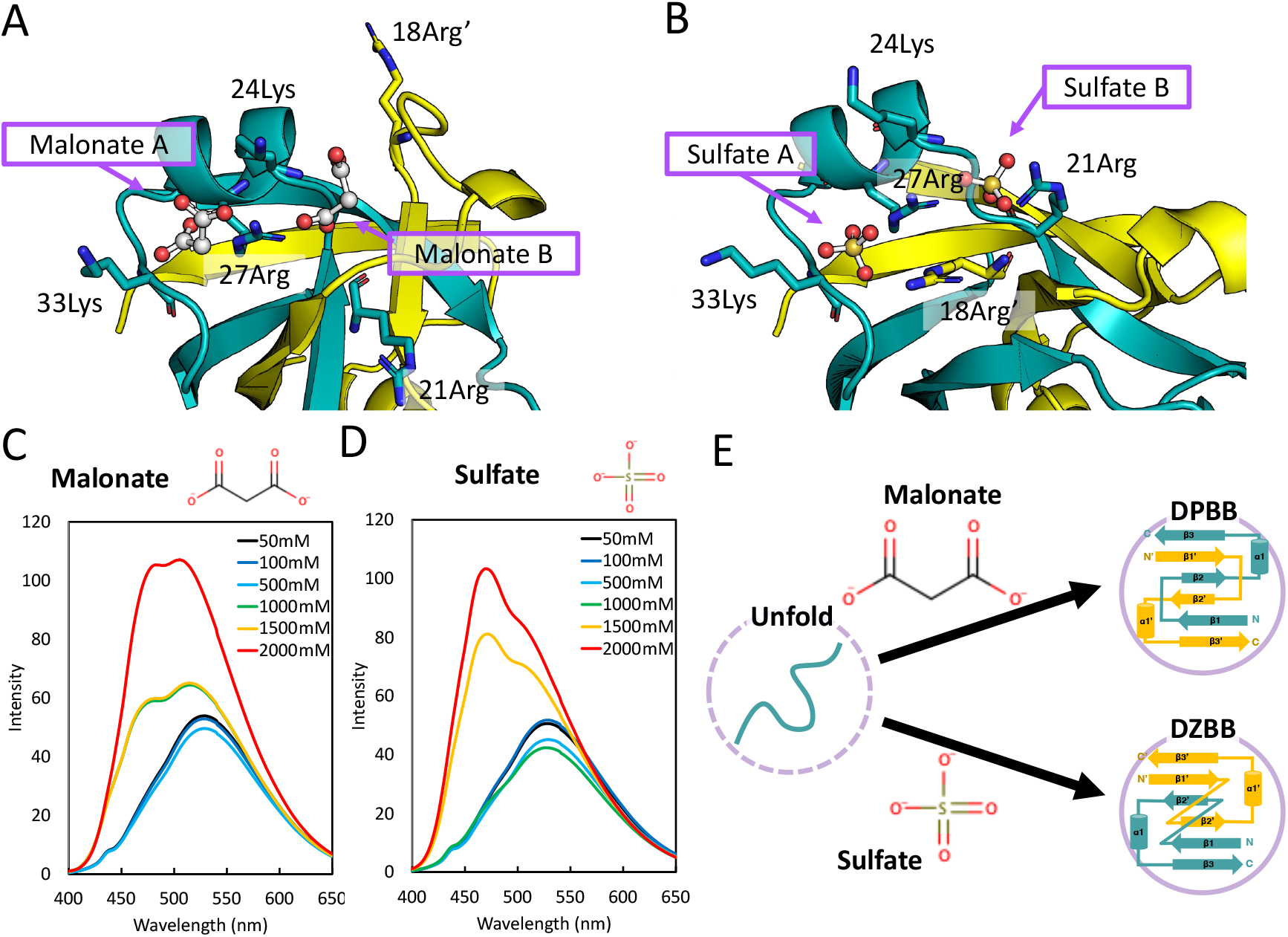
Induction of fold changes by malonate and sulfate ions. (A) Two malonates in the crystal structure of chemically synthesized mk2h_ΔMILPYS in the DPBB form. (B) Two sulfates in the crystal structure of chemically synthesized mk2h_ΔMILPYS in the DZBB form. The five residues related to the coordination of ions are shown as stick models. (C and D) The ANS fluorescence spectra of mk2h_ΔMILPYS in the presence of increasing concentrations of (C) malonate or (D) ammonium sulfate. (E) The scheme of the dual-folding mechanism of mk2h_ΔMILPYS dependent on the types of ions.

To test whether these small molecules can facilitate the folding of mk2h_ΔMILPYS in solution, we analyzed its conformational change by monitoring the binding of the fluorescence probe 8-anilino-1-naphthalenesulfonic acid (ANS). ANS typically binds to a hydrophobic patches of the folding intermediates and molten globules, which changes its fluorescence spectrum. A low concentration of malonates (50–500 mM) did not alter the ANS fluorescence spectra (Fig. 2C). However, the fluorescence signal was increased at over 1,000 mM of malonates, and its peak was blue-shifted (Fig. 2C), implying that high concentrations of malonate induce at least partial folding of mk2h_ΔMILPYS. We also found that high concentrations of sulfate ion (≥ 1,500 mM) increased ANS fluorescence (Fig. 2D). Its spectral pattern was slightly different compared to malonates, perhaps due to the difference in the DPBB- and DZBB-folds. The secondary structure changes depending on the sulfate ions were also observed in the CD analyses (Fig. S8A). These experiments demonstrated that the small ions mediate the folding of mk2h_ΔMILPYS, and their types may lead to two different structures (Fig. 2E).

Interestingly, ANS experiments showed that similar ions, phosphate, malic acid, and citrate, also promoted conformational changes (Supplemental text 2 and Fig. S9). As these anionic ions probably existed in the primordial cells and on the early Earth(*22*–*25*), they might have served as chemical chaperones to enhance the folding of ancient proteins and could have compensated for the low stabilities of evolutionary intermediates during the folding transition (Supplemental text 3, Figs. S7–9).

## The transformation from DZBB to the monomeric OB fold

In the homo-dimeric DPBB and RIFT structures, the β-strands from a monomer are mostly interlaced with the β-strands from the other chain, and thus these peptides can only fold as homo-dimers (Fig. 1B, E). In contrast, in the structure of DZBB, the secondary structure elements from a single subunit are mostly clustered together as in a monomeric protein, except for the swapped β2 strands (Fig 1C, D). Surprisingly, a structural similarity search using the DALI software(*26*) detected high correspondence between the monomeric part of DZBB and OB-fold proteins. In the superimposition of the DZBB and OB proteins, the β1, β2, and β3 strands of DZBB are well aligned with the β1, β3, and β4 strands of the OB-fold protein, respectively (Fig. 3A). In addition, their sequences are partially similar (Fig. S10A). The only significant differences between their structures are the presence and absence of a few secondary structural elements (Fig. 3B). The DZBB monomer lacks two β-strands conserved in the OB-fold (β2 and β5), and while the β2 strand is conserved within OB-fold proteins, the β5 strand is poorly conserved or even absent in some OB proteins; e.g., ribosomal protein L2. Helix α1 of the OB-fold (corresponding to α1 of DZBB) is also missing in some OB proteins (e.g., rL2, S17, and S28). Thus, the original OB-fold has been considered to be a four-stranded β-barrel, of which only β2 is absent in DZBB(*10*).

**Fig. 3.**
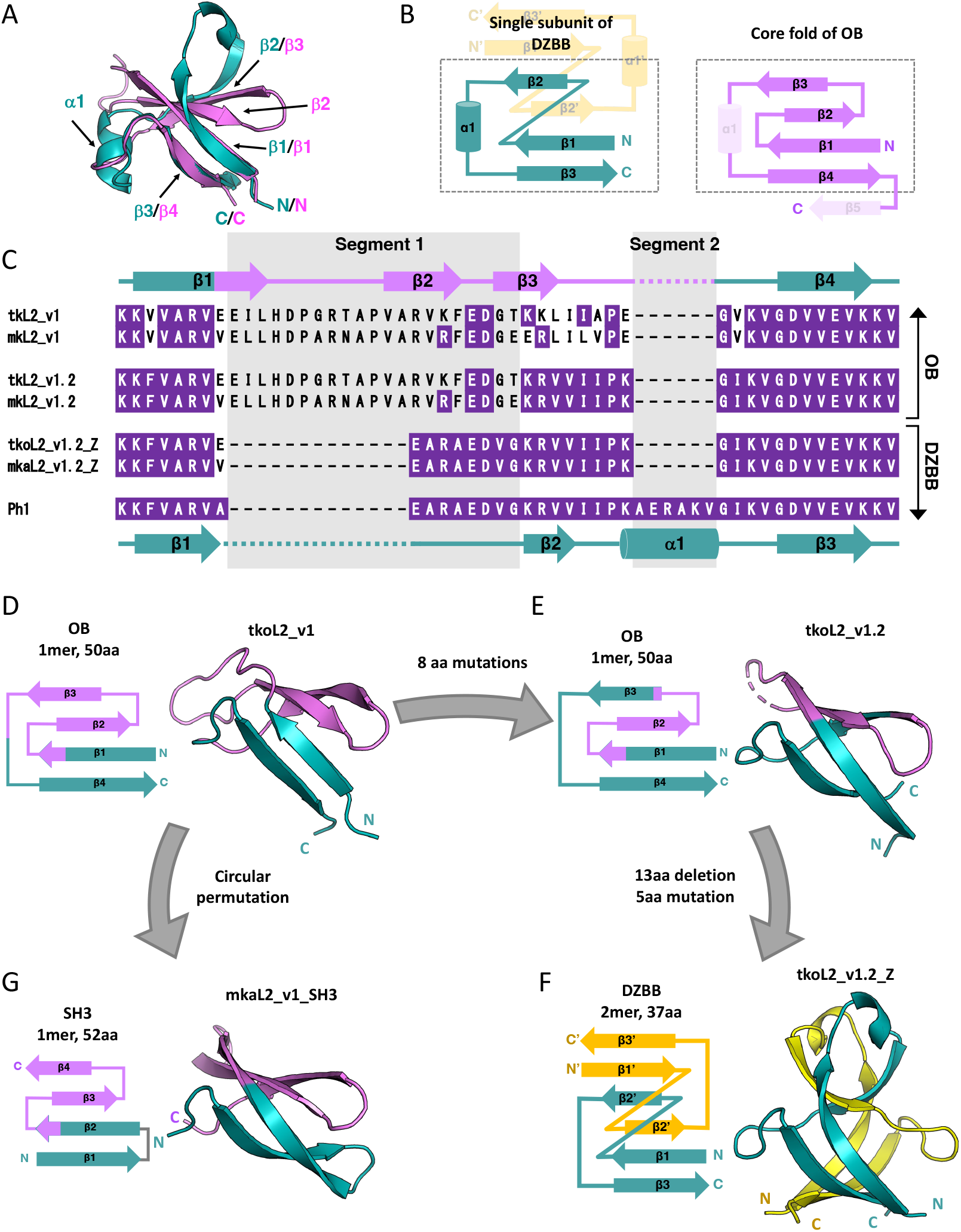
Experimental reconstruction of the evolutionary pathway between the DZBB, OB, and SH3 folds. (A) Superimposed structures of the single subunit of mk2h_ΔMILPYS in the DZBB-fold and the OB domain of the ribosomal protein L2 from *Thermococcus kodakarensis*. (B) Comparison of the topologies of the DZBB and OB folds. In the OB fold, the non-essential secondary elements α1 and β5 are shown as dashed lines. (C) Multiple sequence alignment of the DZBB-OB chimeric proteins with the parent DZBB protein, Ph1. The identical residues with Ph1 are colored purple. The two significantly different segments between tkoL2_v1.2 (OB) and Ph1 (DZBB) are highlighted in gray. (D–G) Crystal structures and topological images of (D) tkoL2_v1, (E) tkoL2_v1.2, (F) tkoL2_v1.2_Z, and (G) mkaL2_v1_SH3. The flows of the engineering procedures are shown as arrows.

To demonstrate the hypothesized interconversion between the DZBB-fold and the OB-fold, we created their chimeric proteins by combining the mk2h_ΔMILPYS and OB proteins. The sequences of the OB domains in the rL2 proteins from thermophilic archaea, *Thermococcus kodakarensis* and *Methanopyrus kandleri*, resemble that of mk2h_ΔMILPYS, while helix α1 is absent in the rL2 proteins. In particular, the OB-domain of rL2 from *M. kandleri* exhibited a high similarity with mk2h_ΔMILPYS (identity 25%, Fig. S10A). Given that the mk2h_ΔMILPYS was originally constructed from the DPBB protein from *M. kandleri*,(*7*) the genome of this archaeon might still preserve the evolutionary information of ancient proteins. The sequence region surrounding β2-β3 of the OB-domain in rL2 was incorporated into the corresponding position of mk2h_ΔMILPYS, based on the superimposed structures (Fig. 3A). Through this process, we constructed six unique variations by modifying the positions and extents of insertion (Fig. S10B and Table S1). Biochemical and crystallographic analyses revealed that two of the six chimera proteins, tkoL2_v1 and mkaL2_v1 (Fig. 3C), adopt stable four-stranded OB-folds (Figs. 3D, S5C and S11). The DZBB and OB fragments comprise approximately 40 and 60% of these chimeric proteins, respectively. Therefore, the monomeric OB-fold could be reconstructed by simply combining the DZBB and OB-fold protein segments, without optimizing the structural interfaces between both parts.

To examine what determines the DZBB–OB transition, additional intermediates have been engineered by sequential mutagenesis and experimental validation steps (Supplemental text 4, Fig. S5D, S12A, and Table S1). The 2nd generation mutants, tkoL2_v1.2 and mkaL2_v1.2, have ∼60% sequence identities with Ph1 (DZBB-fold), and retained moderate thermostability (Figs. 3C and S13). In these designs, only a short segment was from rL2 (segment 1: from the middle of β1 to the start of β3) and the other parts are from Ph1 (Figs. 3C and S12A). Helix α1 (segment 2) was also omitted. The crystallographic analysis revealed that tkoL2_v1.2 still adopts the four-stranded OB fold (Fig. 3E), indicating that segment 1, but not segment 2, could serve as the determinant for the fold transition between the DZBB and OB structures. To test this assumption, we conducted the reverse engineering by replacing segment 1 of tkoL2_v1.2 and mkaL2_v1.2 with the 8-amino acid sequence forming the Z-loop of Ph1 (the 3rd generation mutants: tkoL2_v1.2_Z and mkaL2_v1.2_Z, Figs. 3C, S12B, Table S1). SEC and CD experiments demonstrated that both mutants folded with β-structures and had moderate thermal stabilities (Fig. S14). The crystal structures of tkoL2_v1.2_Z and mkaL2_v1.2_Z revealed that they adopt the DZBB fold even without the α1 region (segment 2) (Figs. 3F and S5E). These results demonstrated that the segment 1 is sufficient to archive the fold-change from DZBB to OB.

Furthermore, we verified that the 13 a.a. sequence forming the flexible β-turn in segment 1 in the OB-fold designs could be shortened to 7 a.a. (Supplemental text 4, Figs. S15–17, and Table S1). Taken together, very short In/Del and a few point mutations are the determinants in the transition between the DZBB and OB folds. This facile interchangeability between DZBB and OB folds suggests that such a drastic fold transition likely occurred in the early evolutionary stage of life.

## Transformation from OB to SH3

Given the high similarity between the OB and SH3 folds, they are presumed to have evolved from a common ancestral protein (Fig. S1)(*9, 10*). Loren Williams’ group suggested that the four-stranded core fold of OB could have transformed to SH3 by a simple circular permutation (or vice versa)(*10*). Following this evolutionary hypothesis, we tried to convert the reconstructed OB proteins to the SH3 fold. The fourth strand of tkoL2_v1 and mkaL2_v1 was trimmed and connected to the N-terminal end by two residues “DG,” an ideal sequence to form a β-turn(*19*) (tkoL2_v1_SH3 and mkaL2_v1_SH3)(Table S1). While tkoL2_v1_SH3 had a random-coil structure, mkaL2_v1_SH3 exhibited a characteristic CD spectrum for a β-sheet protein and remained almost intact even at 90°C (Fig. S18). We also determined its crystal structure and confirmed that mkaL2_v1_SH3 adopted the four-stranded SH3 fold (Fig. 3G). This experimental conversion from OB to SH3 strongly supports the previous hypothesis that these folds could have emerged by a simple permutation of their four-stranded core fold(*10*).

## DNA binding abilities of the reconstructed β-barrels

As most β-barrels in the central dogma machinery function by interacting with nucleic acids, we investigated the DNA or RNA binding capabilities of the reconstructed β-barrels by an electrophoresis mobility shift assay (EMSA) (Fig. 4 and Fig. S19). When mixing the stable DPBB protein (mk2h_ΔMILPS) and the 20 bp double-stranded DNA (dsDNA), some portion of dsDNA was slowed and stacked in the well as previously reported(*7*). Most DNA molecules did not migrate from the well when mixed with the proteins with the stable DZBB fold (Ph1, tkoL2_v1.2_Z, and mkaL2_v1.2_Z) and RIFT fold (Ph1_DG and Ph1_GG), indicating that these proteins formed large aggregates with dsDNA (Fig. 4A). In particular, Ph1, mkaL2_v1.2_Z, Ph1_DG, and Ph1_GG interacted with dsDNA even in high salt conditions (500 mM NaCl) (Fig. S19B). No significant sequence specificity was observed when we tested another 20 bp dsDNA (Fig. S19C). These proteins might interact with the phosphate groups in the DNA backbone in a similar way to the sulfate ions in the crystal structures of DZBB (Fig. 2B). The proteins with the DPBB, DZBB, and RIFT folds also interacted with ssDNA and ssRNA (Fig. S19D and E). These findings suggest that, like their modern descendants, the ancient DPBB, DZBB, and RIFT proteins also interacted with nucleic acid polymers.

**Fig. 4.**
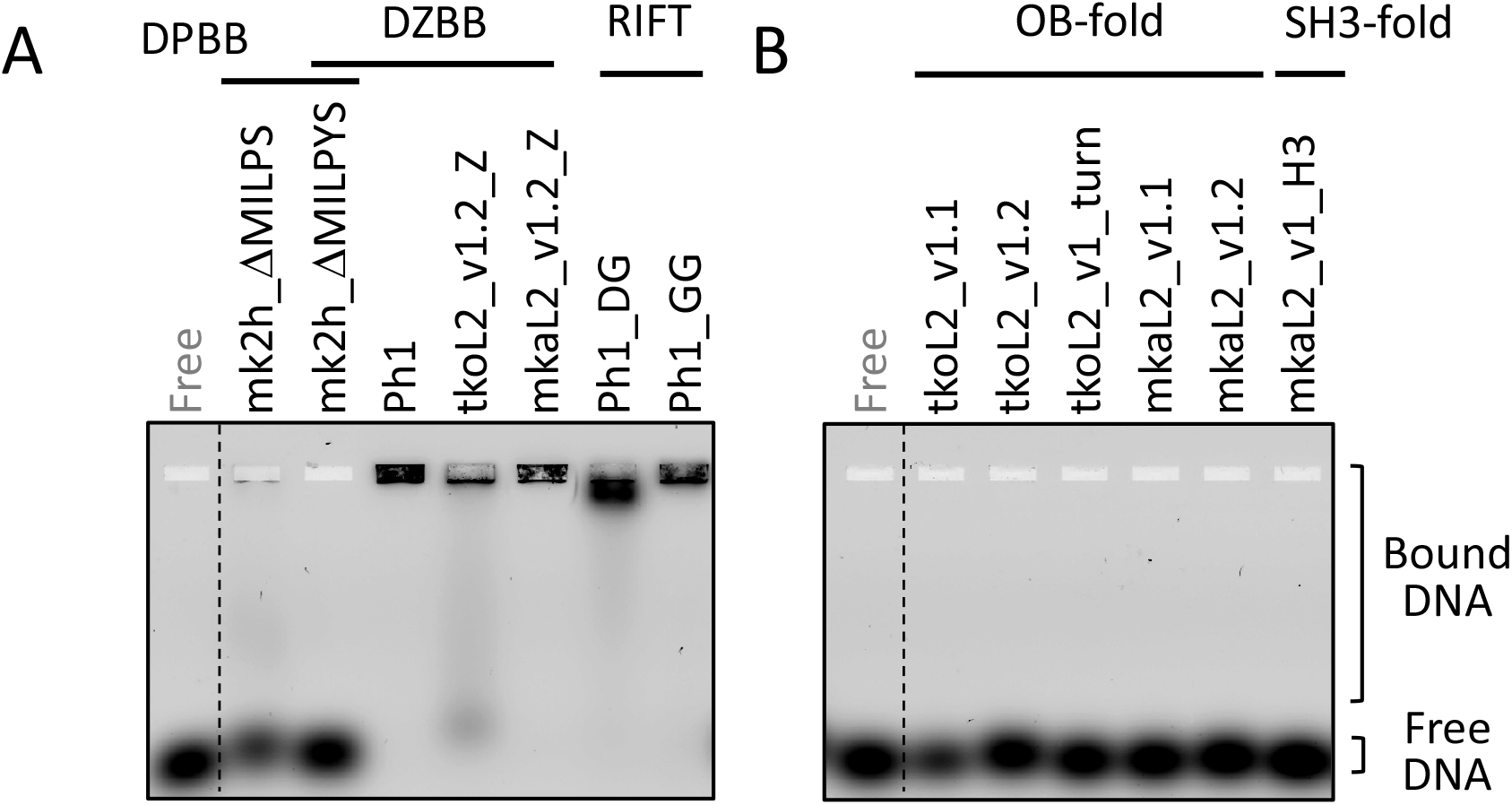
Electrophoresis mobility shift assay to analyze the dsDNA binding properties of the reconstructed β-barrel proteins. (A) 10 nM FAM-labeled dsDNA was mixed with 5 μM proteins with the DPBB, DZBB, and RIFT folds, and then the mixtures were subjected to 2% agarose gel electrophoresis. (B) The EMSA results for the proteins with the OB and SH3 folds.

In contrast, the reconstructed OB and SH3 proteins did not interact significantly with any oligonucleotides (Fig. 4B and S19F–H). Only high concentrations of tkoL2_v1.2 and mkaL2_v.1.2 retarded the migration of dsDNA slightly (Fig. S19I). The acquisition of the weak DNA binding affinity of these two proteins may have resulted from additional lysine residues at the end of β3 and the 3rd loop, compared to other OB-fold chimeras (Fig. 3C). While the typical modern OB-fold proteins interact with an oligonucleotide at the surface of β1–3 (β2–4 in the SH3-fold proteins)(*27*), the corresponding region of ribosomal protein L2 used in this report (β2–3) does not directly interact with rRNA in the ribosome, likely resulting in the weak affinity of the reconstructed OB and SH3 proteins.

## Conclusion

In this report, we demonstrated that the short and simple peptide, mk2h_ΔMILPYS, adopts to not only the homo-dimeric DPBB fold but also another novel homo-dimeric β-barrel fold, DZBB, as a metamorphic protein (Figs. 1 and 2). In addition, from the DZBB fold, the evolutionary pathway among distinct β-barrel folds, RIFT and OB, could be reconstructed by simple and feasible mutation steps. A single deletion in the DZBB sequence converted it into the RIFT fold (Fig. 1). In contrast, the single insertion of a short sequence forming a β-strand and a few point mutations converted it to an OB fold, accompanied by the oligomeric state change from dimer to monomer (Fig. 3). Furthermore, the reconstructed OB fold could also be converted to the four-stranded SH3 fold by a simple circular permutation (Fig. 3E). Thus, these ancient β-barrel folds (DPBB, RIFT, OB, and SH3) could be readily interconverted by a few feasible mutations through the potential missing link, DZBB (Fig. 5 and Supplemental text 5), indicating their significantly close evolutionary relationships (Supplemental text 6).

**Fig. 5.**
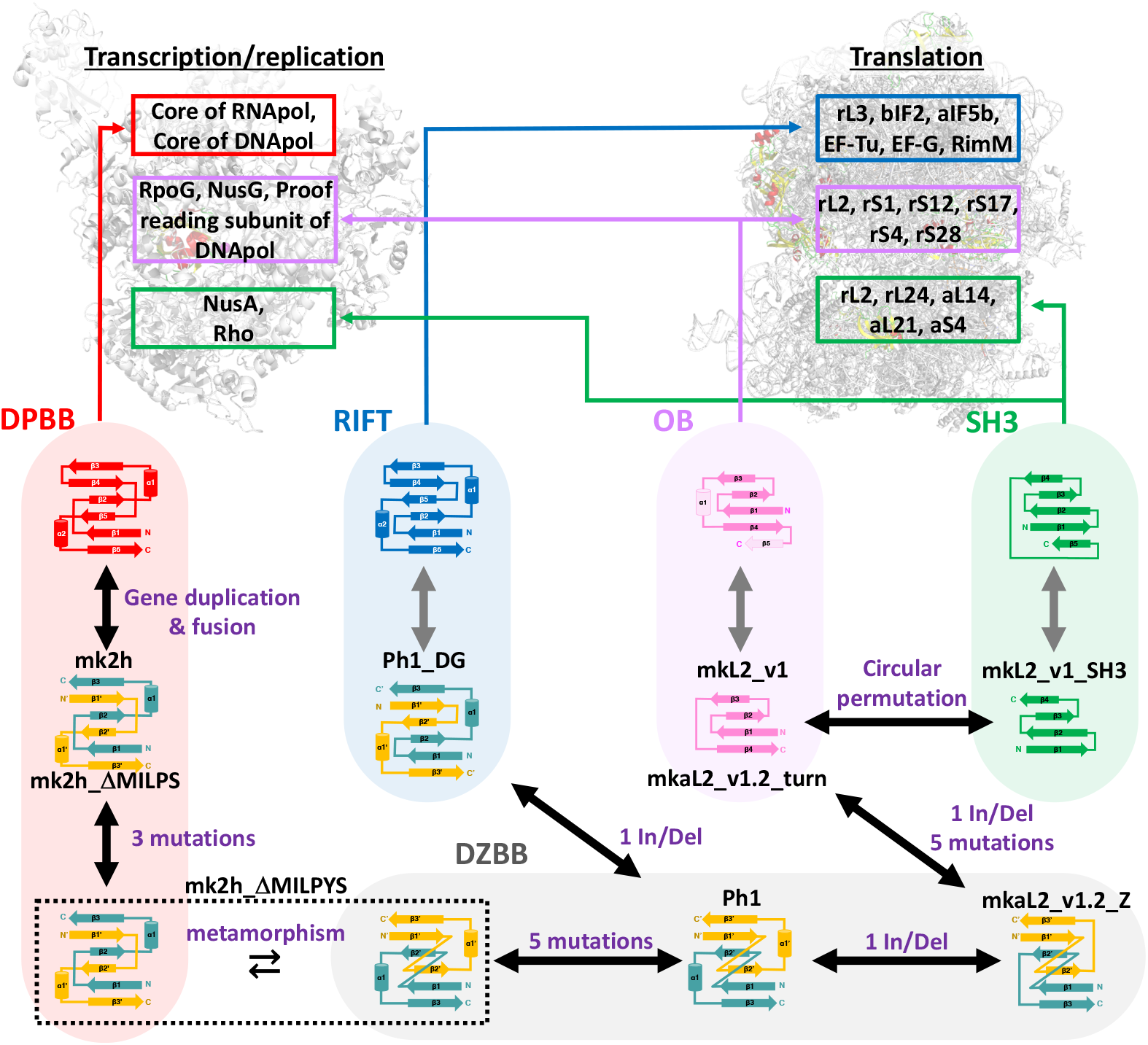
The proposed evolutionary network between the modern four β-barrels through the missing link fold, DZBB. The experimentally verified routes between each mutant pair are connected with black arrows, and the minimal necessary mutations between designs are written in purple. The DPBB protein’s evolutionary pathway, which was examined in the previous study, is highlighted in a light red background. The evolutionary routes of DZBB, RIFT, OB, and SH3 are highlighted in gray, light blue, pink, and green backgrounds, respectively. Representative proteins conserving each barrel fold are shown in the upper panel.

This also implies that the diverse β-barrel folds can be produced from a limited sequence space. In an evolutionary time scale, the variety of ancient β-barrel folds might have diverged in a very short period, like the rapid diversification of animal species in the Cambrian. This rapid diversification of the various β-barrel folds probably preceded and primed the subsequent development of the elaborate molecular machines underlying the central dogma(*28*–*32*) (Fig. 5). Moreover, the reconstructed β-barrels (except for the mutant with the SH3 fold) retained the DNA and RNA binding affinities (Figs. 4 and S19). Thus, during the diversification process of these folds, the fundamental nucleic-acid-binding property might have been inherited by the daughter folds, which then became specialized to the specific substrates and enzymatic reactions by stepwise mutations in each lineage. The evolutionary pathways depicted here provide the groundwork for more detailed and broader studies of early protein evolution and the origin of the central dogma.

## Acknowledgments

This work is based on experiments performed at KEK (project number: 2020G056 and 2022G005), SPring-8, and SLS. The authors are grateful to the beamline staff scientists at KEK, SPring-8, and SLS. We thank Hideaki Niwa, Toshiaki Hosaka, and Kentaro Ihara for assistance with the X-ray diffraction experiments. We also thank Hongding Liu for assistance of the DNA cloning experiment. S.Y. and S.T. were supported by JSPS (18H01328, 20K15854, and 22H01346). S.T. was also supported by the Astrobiology Center Program of National Institutes of Natural Sciences (AB0503).

